# The adaptor protein NumbL is involved in the control of glucolipotoxicity-induced pancreatic beta cell apoptosis

**DOI:** 10.1101/2020.08.16.253286

**Authors:** Halesha D. Basavarajappa, Jose M. Irimia, Patrick T. Fueger

## Abstract

Avoiding loss of functional beta cell mass is critical for preventing or treating diabetes. Currently, the molecular mechanisms underlying beta cell death are partially understood, and there is a need to identify new targets for developing novel therapeutics to treat diabetes. Previously, our group established that Mig6, an inhibitor of EGF signaling, mediates beta cell death under diabetogenic conditions. The objective of this study was to clarify the mechanisms linking diabetogenic stimuli to beta cell death by investigating Mig6-interacting proteins. Using co-immunoprecipitation and mass spectrometry, we evaluated the binding partners of Mig6 under both normal glucose (NG) and glucolipotoxic (GLT) conditions in beta cells. We identified that Mig6 interacts dynamically with NumbL; whereas Mig6 associates with NumbL under NG, this interaction is disrupted under GLT conditions. Further, we demonstrate that siRNA-mediated suppression of NumbL expression in beta cells prevented apoptosis under GLT conditions by blocking activation of NF-κB signaling. Using co-immunoprecipitation experiments we observed that NumbL’s interactions with TRAF6, a key component of NFκB signaling, are increased under GLT conditions. The interactions among Mig6, NumbL, and TRAF6 are dynamic and context-dependent. We propose a model wherein these interactions activate pro-apoptotic NF-κB signaling while blocking pro-survival EGF signaling under diabetogenic conditions, leading to beta cell apoptosis. These findings indicate that NumbL should be further investigated as a candidate anti-diabetic therapeutic target.

## 1. Introduction

Diabetes is a complex metabolic disorder affecting nearly 463 million people worldwide according to the International Diabetes Federation. The disease is characterized by increased blood glucose levels due to an imbalance in whole-body glucose homeostasis. Insulin-secreting pancreatic beta cells play a key role in controlling glucose homeostasis, and the depletion of functional beta cell mass is central to the manifestation of diabetes [1]. Hence, preventing beta cell loss or restoring functional beta cell mass remain major challenges in finding cures for diabetes.

Functional beta cell mass is modulated by multiple processes such as beta cell proliferation, beta cell hypertrophy, insulin secretory capacity, and apoptosis. During the development of type 2 diabetes (T2D), initially pancreatic beta cell mass increases to maintain glucose homeostasis under chronic nutrient oversupply and peripheral insulin resistance conditions [2]. This increase in beta cell mass compensates for the increased insulin demand under excess glucose levels and is partially driven by low levels of ER stress through unfolded protein response [3]. However, if the secretory burden persists, unresolved ER stress leads to beta cell death [4, 5]. Moreover, the glucolipotoxic (GLT) milieu (elevated levels of both glucose and lipids), prevalent in diabetic conditions, further deteriorates the survival capacity of beta cells [6–9]. Once functional beta cell mass declines, overt T2D arises [10]. Understanding the molecular events leading to pancreatic beta cell death under diabetogenic conditions is essential for discovering new therapies to prevent or cure diabetes. Apoptosis is a major pathway for the beta cell death [11] and is induced by absence of pro-survival signals such as EGF signaling [12] and activation of proapoptotic pathways such as NF-κB signaling [13].

It is well established that GLT induces NF-κB signaling, and this pathway induces beta cell apoptosis [13, 14]. Inactive NF-κB is bound to an inhibitor protein, IκB, and sequestered in the cytosol. The NF-κB pathway is activated when IκBα is phosphorylated and degraded to release free NF-κB. Once free of inhibition, NF-κB is phosphorylated and migrates to the nucleus to regulate transcription [15]. This degradation of IκBα is initiated by a sequence of events involving IKK, NEMO and TRAF6 [15, 16]. TRAF6, upon activation, is polyubiquitinated and then activates NEMO and IKK, which in turn phosphorylate IκBα [16]. However, how NF-κB signaling is activated under GLT is not clearly understood.

In addition to the induction of proapoptotic pathways, the absence of pro-survival signals also contributes to beta cell apoptosis. Among the many proliferative pathways characterized, epidermal growth factor receptor (EGFR) signaling is crucial for both beta cell proliferation and survival [12, 17], and EGF treatment has been reported to induce beta cell proliferation and restore beta cell mass in rodents [18]. However, EGF signaling is impaired during beta cell stress, as occurs in T2D [19].

Previously, our group has demonstrated that Mig6, an inhibitor of EGFR signaling, plays a key role in beta cell apoptosis under diabetogenic conditions [19–21]. Upon induction, Mig6 binds to the EGF receptor and blocks pro-proliferative signaling [22]. Moreover, we have observed that under GLT conditions, Mig6 levels are elevated, and EGFR signaling is inhibited (Y-C Chen and PT Fueger, personal communications). Although Mig6 is considered a molecular brake for proliferation, it also antagonizes beta cell survival and impairs beta cell function [19, 20]. Thus, strategies for suppressing Mig6 action in beta cells could improve the retention of functional beta cell mass.

The objective of this study was to better understand how GLT regulates beta cell death by investigating Mig6 binding partners. We discovered that Mig6 dynamically interacts with the adaptor protein NumbL, a component of Notch signaling [23, 24]. Furthermore, we delineated a novel role for NumbL in beta cell apoptosis, and we propose that reducing NumbL could be exploited to prevent beta cell death under glucolipotoxic conditions.

## 2. Results

### 2.1 Mig6 dynamically interacts with NumbL protein

Although Mig6 was initially considered as a stress-inducible gene, increasing reports indicate that it contributes to beta cell death under stressed conditions [19, 20, 30]. To clarify the mechanisms underlying diabetogenic beta cell death, we sought to identify Mig6 associating partners in a GLT cell culture model, which mimics the diabetic milieu [31, 32]. To identify potential interacting protein partners of Mig6, we overexpressed FLAG-tagged Mig6 protein in Ins-1-derived 832/13 cells using adenovirus and exposed the cells to NG or GLT conditions for 12 h. Using an anti-FLAG antibody, we immunoprecipitated Mig6 along with its interacting proteins and identified the associating proteins using mass spectrometry (**Figure 1A**). As Mig6 is an adaptor protein, it was expected that multiple proteins bind to Mig6 (**Figure 1A**). Hence, we screened for interacting proteins whose binding pattern with Mig6 was significantly altered with changes in the glucolipotoxic milieu of the culture conditions (**Figure 1B**). Additionally, protein components of the cytoskeleton system such as actin, spectrin, and myosin would be ignored. Given the goals of the screen, we focused on proteins reported to be involved in EGF and/or NF-κB signaling. We then confirmed the results by immunoblot analysis and reverse co-immunoprecipitation experiments (**Figure 1C-D**). We observed that Mig6 interacted dynamically with NumbL protein, showing strong association under NG conditions and weaker association under GLT conditions (**Figure 1C-D**).

**Figure 1:**
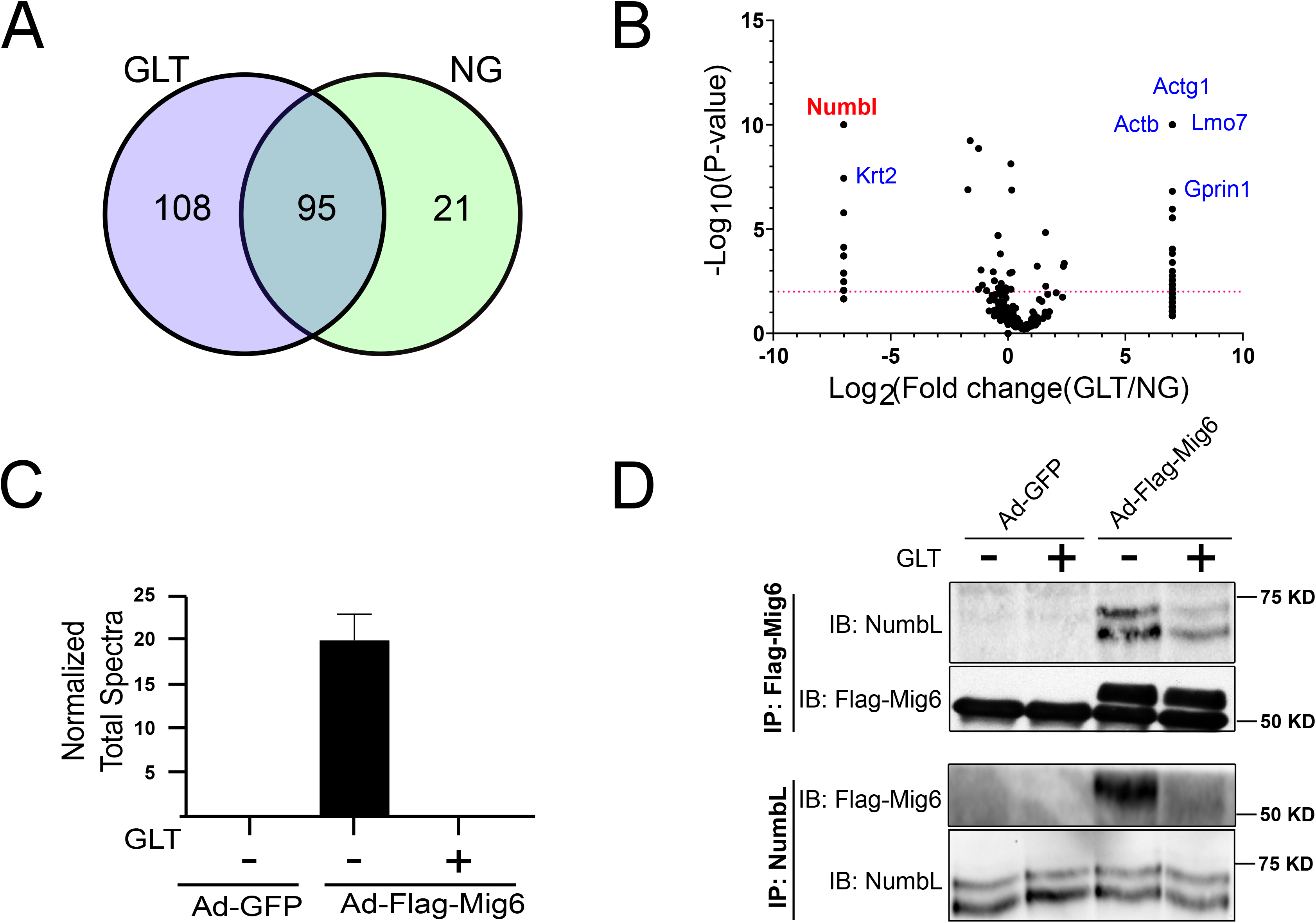
Mig6 interacts with NumbL under NG and this interaction is disrupted under GLT conditions: (A) Venn diagram depicting number of proteins pulled down with Flag-Mig6 under NG and GLT conditions. (B) The fold change in number of peptide counts of each pulled down protein under GLT and NG conditions is plotted against P-values of Fisher’s exact *t*-test to screen for proteins with significantly altered binding pattern with Mig6. (C) the number of unique peptide counts of NumbL protein identified from eluate samples (normalized to total spectral count of the eluate samples). (D) Representative immunoblot from co-immunoprecipitation experiments confirming the interaction between NumbL and Mig6 under NG and GLT conditions. *GLT*, glucolipotoxicity; *NG*, normal glucose.

To verify that the decreased binding was not due to decreased expression levels of NumbL, we measured both the mRNA and protein levels of NumbL under both NG and GLT conditions at various time points (**Figure 2**). We detected no change in the levels of NumbL in 832/13 cells exposed to GLT conditions, suggesting that the interaction between Mig6 and NumbL is dynamic and not determined solely by protein abundance. Similarly, primary human islets cultured in GLT conditions for 48 h exhibited no change in NumbL expression levels (**Figure 2D**). Further, we measured NumbL mRNA levels in primary islets from patients with pre-diabetes (HbA1c level between 5.7-6.4%) and T2D (HbA1C levels > 6.4%) (**Figure 2C**). Although there were no significant changes in expression levels between controls (HbA1C levels < 5.7%) and patients with diabetes, surprisingly there was a significant decrease in NumbL between the control and pre-diabetic groups. Further investigation is required to understand this difference in the NumbL levels between control and pre-diabetic subjects.

**Figure 2:**
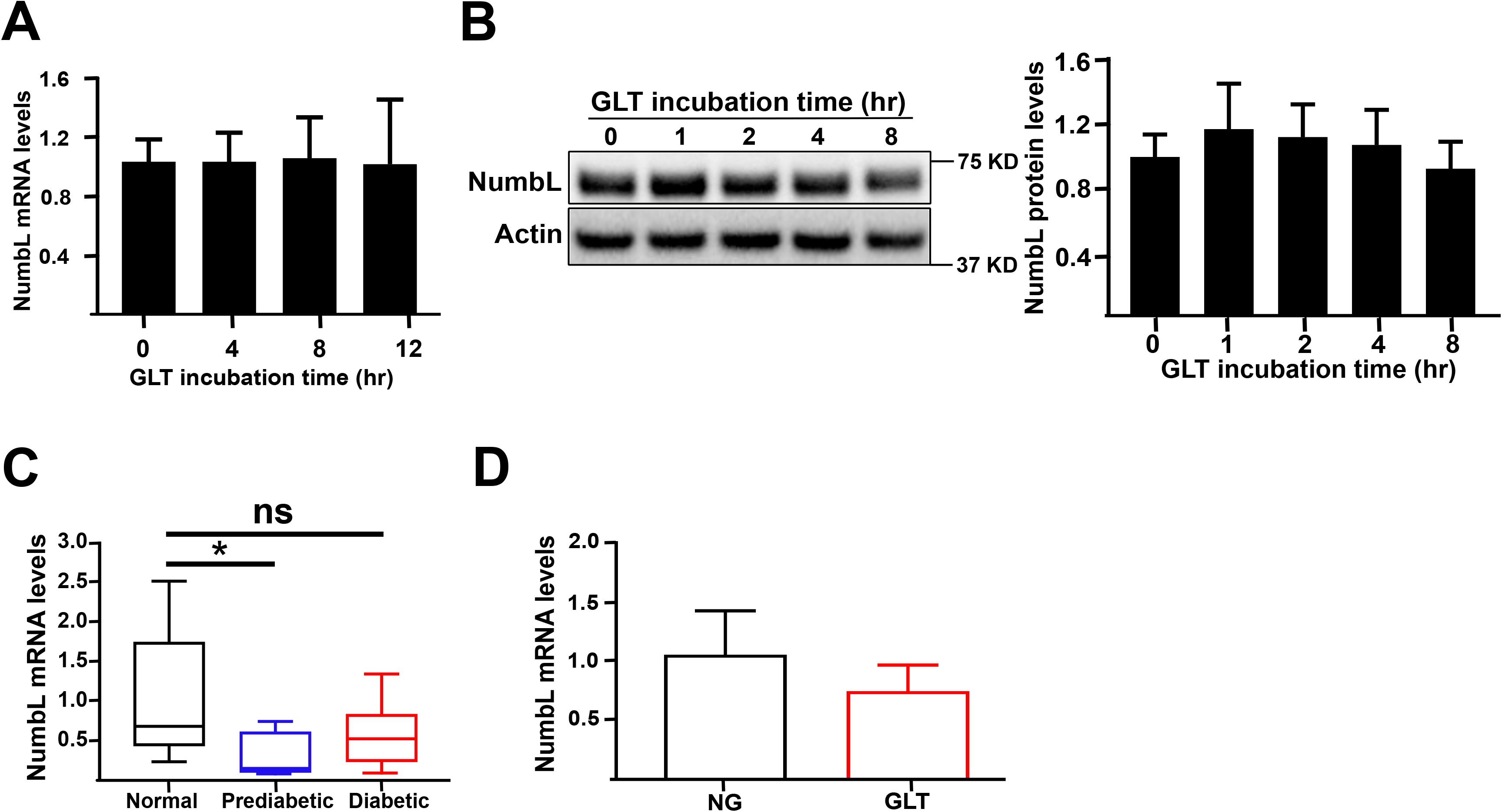
NumbL expression levels are unchanged in beta cells under GLT conditions: (A) mRNA levels of NumbL in 832/13 cells after culture in GLT conditions for the indicated time periods. (B) Protein levels of NumbL in 832/13 cells after culture in GLT conditions for indicated time periods. (C) mRNA levels of NumbL in human islets obtained from normal, prediabetic and diabetic patients. * p<0.05 vs. normal. (D) mRNA levels of NumbL in human islets after culture in NG and GLT conditions for 48 h. All data are represented as Mean +/- SEM and were compared by ANOVA.

### 2.2 Down regulation of NumbL prevents glucolipotoxicity-induced beta cell apoptosis

As Mig6 has been implicated in inducing beta cell apoptosis under stressed conditions, we investigated the role of its interacting protein, NumbL, in beta cell apoptosis under GLT conditions. As depicted in **Figure 3A**, GLT induced apoptosis in 832/13 cells in a time-dependent manner, as measured by cleaved caspase 3 activity. To test the role of NumbL, we used siRNA knockdown and confirmed over 80% suppression of NumbL expression by immunoblot analysis (**Figure 3B**). The activation of apoptosis with GLT was significantly reduced by siRNA-mediate suppression of NumbL as measured by the levels of cleaved caspase 3 (**Figure 3C**) and cleaved caspase-3 activity (**Figure D**).

**Figure 3:**
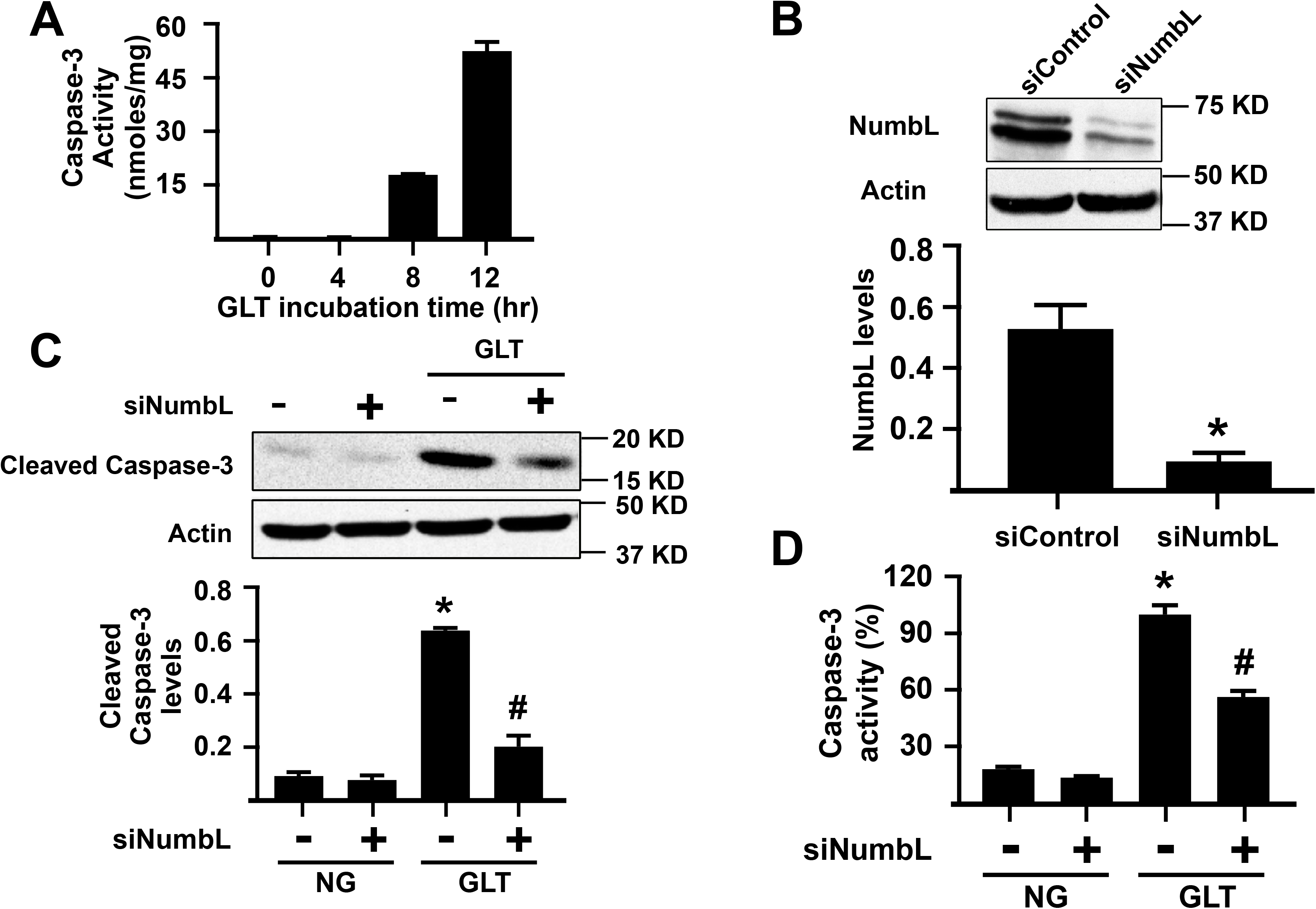
siRNA-mediated knock down of NumbL inhibits GLT-induced apoptosis of Ins1 832/13 cells. (A) Cleaved caspase 3 activity after exposure of cells to GLT for the indicated time periods. (B) Representative blot and quantification of NumbL protein levels from cells treated with a control RNAi (siControl) or RNAi to NumbL (siNumbL). * P<0.05 (*t*-test). (C) Representative blot and quantification of cleaved caspase-3 protein from cells treated with siNumbL either under NG or GLT for 12 h. *p<0.05 vs. siControl treated under NG; #p<0.05 vs. siControl treated under GLT (ANOVA). (D) Cleaved caspase-3 activity in siRNA-treated cells after culture in NG and GLT conditions for 12 h. *p<0.05 vs. siControl treated with NG; #p<0.05 vs. siControl treated with GLT. The data are representative from at least three independent experiments.

The prevention of beta cell apoptosis under GLT prompted us to test the extent to which NumbL down regulation resolved ER stress induced by GLT. However, there was no significant change in GLT-induced ER stress between the siRNA control and siNumbL groups, as measured by the levels of phospho-eIF2a (**Supporting Information Figure S1**).

### 2.3 Reducing NumbL does not enhance beta cell proliferation and insulin secretion

In lung cancer cell lines, suppression of NumbL has been reported to promote cell proliferation [33]. Hence, we sought to determine the extent to which NumbL knock down in 832/13 cells induces beta cell proliferation using an EdU incorporation assay. However, NumbL knock down did not induce significant proliferation as compared to control siRNA-treated cells under either NG or GLT conditions (**Supporting Information Figure S2A**). This result is further corroborated by the observation that knock down of NumbL did not rescue the glucolipotoxicity-induced impairment of EGF signaling, a key pro-proliferative pathway in beta cells (**Supporting Information Figure 2B**). However, it is interesting to note that NumbL knock down significantly increased EGFR phosphorylation under NG conditions. Further, NumbL depletion did not affect beta cell function, as measured by glucose-stimulated insulin secretion (**Supporting Information Figure S2C**).

### 2.4 NumbL reduction does not activate Notch signaling in 832/13 cells

NumbL, along with its homologous protein Numb, were originally characterized as Notch signaling inhibitors and cell-fate-determining proteins [23, 24, 34]. Hence, we tested whether knock down of NumbL activates Notch signaling in response to GLT conditions by monitoring the levels of Hes-1, a downstream target gene of Notch signaling. Whereas Hes-1 expression was increased under GLT conditions in 832/13 cells, knock-down of NumbL did not affect Notch signaling (**Supporting Information Figure S3A**). Interestingly, we noticed that only the siRNA directed against Numb and not NumbL increased Hes-1 expression levels under GLT conditions. These observations indicate that under the studied conditions NumbL may not participate in Notch signaling in 832/13 cells or that Numb is the major mediator of Notch inhibition in 832/13 cells. These observations contrast with the functions of NumbL observed in other cell types [24] indicating tissue-specific variations in the functions of NumbL.

### 2.5 Down regulation of NumbL prevents glucolipotoxicity-induced activation of the NF-κB pathway

Several groups have reported that NF-κB pathway activation leads to apoptosis in beta cells under diabetogenic conditions [13, 35, 36]. In its inactive state, NF-κB is bound to the IκB inhibitor protein and sequestered in the cytosol. Upon activation, IκBα is phosphorylated and degraded by the proteasome. Once free of inhibition, NF-κB is phosphorylated and migrates to the nucleus where it exerts its transcriptional function.

When either human islets (**Figure 4A-C**) or 832/13 cells (**Figure 4D-G**) were incubated in GLT conditions, we observed significant activation of the NF-κB pathway as measured by decreasing levels of IκB and increased phosphorylation of p65, which is activated NF-κB. However, depletion of NumbL in beta cells (**Figure 4D-G**) significantly prevented degradation of IκBα and phosphorylation of p65, indicating that inactivation of NumbL in beta cells can increase resistance to GLT-induced NF-κB pathway activation. These data confirm that NumbL mediates GLT-induced NF-κB pathway activation.

**Figure 4:**
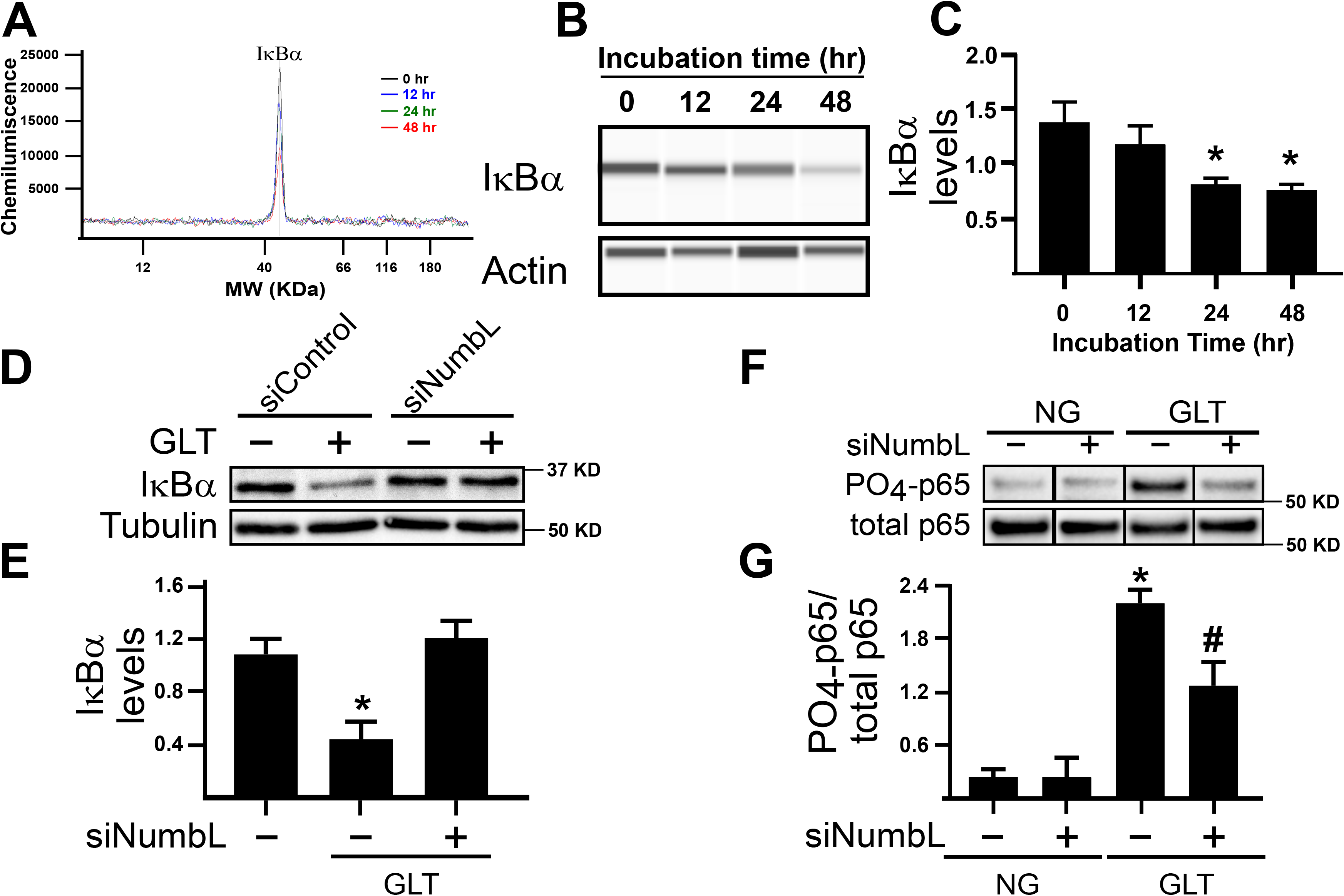
NumbL downregulation prevents GLT-mediated activation of NF-κB signaling. (A) & (B) Representative histogram and immunoblot of IkBa protein using the capillary Western-blot system from human islets treated with GLT conditions for the indicated time periods. (C) Quantification of IkBa protein. The quantification data are from three independent experiments. *p<0.05 vs. ‘0 hr GLT’ group. (D) & (E) Representative immunoblot and quantification of IkBa protein from siRNA-treated 832/13 Ins1 cells cultured in NG or GLT conditions for 12 h. *p<0.05 vs. siControl group under NG. (F) & (G) Representative immunoblot and quantification of phospho-p65 protein from siRNA treated 832/13 Ins1 cells cultured in NG or GLT conditions for 12 hours. *p<0.05 vs. siControl group under NG; #p<0.05 vs. siControl group under GLT.

### 2.6 NumbL interacts with TRAF6 in a context dependent manner

As NumbL down regulation inhibits NF-κB signaling under GLT, we evaluated the mechanism through which NumbL is linked to NF-κB. Several groups have reported that NumbL interacts with TRAF6 and modulates NF-κB signaling [37, 38]. Thus, we tested the dynamics of interactions between NumbL and TRAF6 under GLT in beta cells. As reported in **Figure 5A**, in beta cells, endogenous NumbL and TRAF6 interact under NG conditions and this interaction is increased nearly 2-fold under GLT conditions. To determine if this increased binding is simply due to increased expression of TRAF6, we measured TRAF6 protein levels under GLT over different time points (**Figure 5B**) and noted no significant increase in the TRAF6 expression levels, indicating an increased interaction between NumbL and TRAF6 under GLT. In order to further test the role of TRAF6 in activation of NF-κB signaling and apoptosis under GLT, we treated beta cells under GLT with different concentration of compound 6877002, an inhibitor of TRAF6 activation [39], and monitored the levels of IkBa levels (**Figure 5C**) and cleaved caspase-3 (**Figure 5D**). We observed that compound 6877002 decreased both NF-κB activation and beta cell apoptosis in a dose dependent manner.

**Figure 5:**
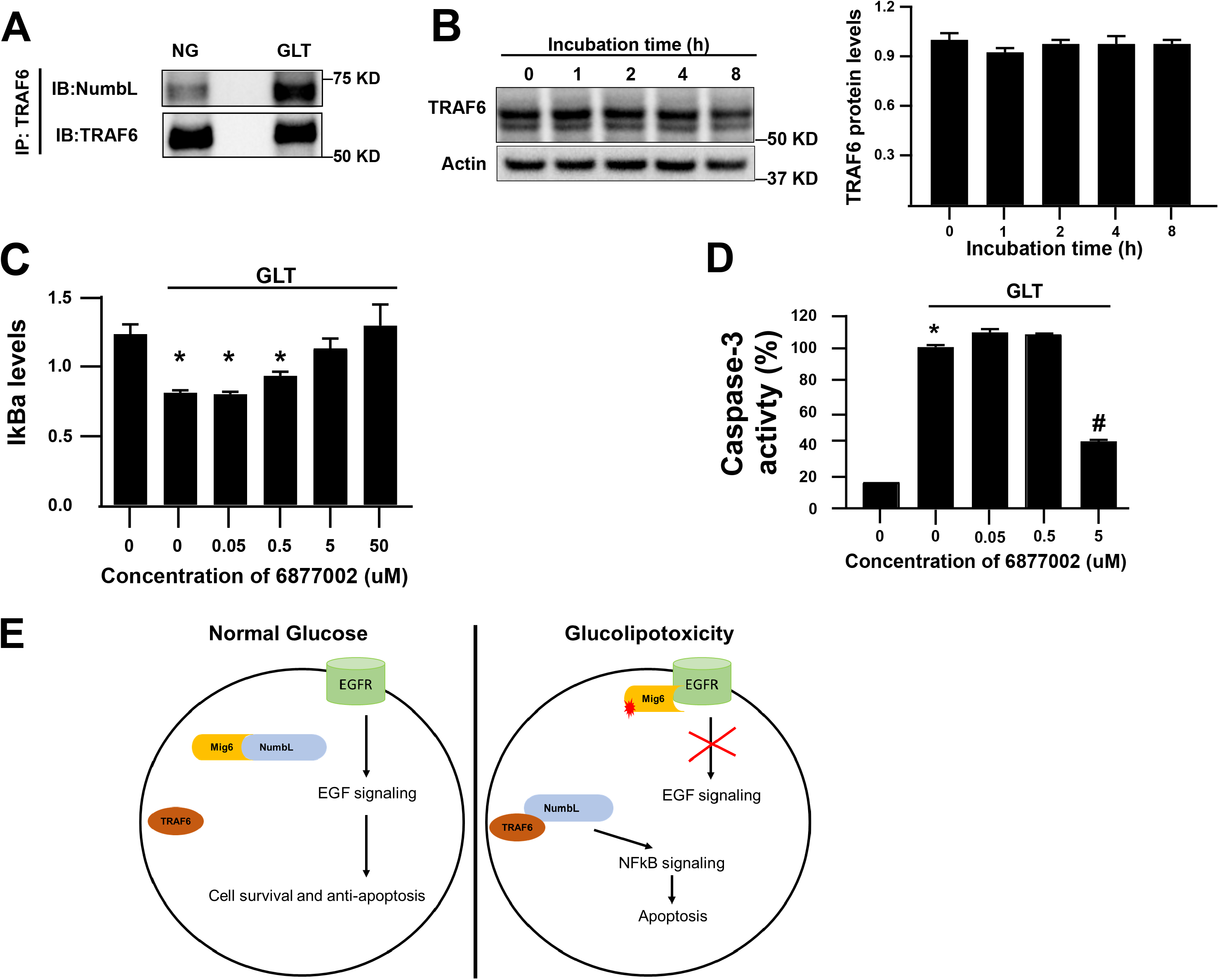
NumbL interaction with TRAF6 is increased under GLT conditions. (A) Representative blot of TRAF6 proteins pulled down using an anti-NumbL antibody. (B) Representative blot of TRAF6 protein and quantification of TRAF6 protein levels in cells treated with GLT for the indicated time periods. (C & D) Effect of TRAF6 inhibitor 6877002 on the levels of IkBa and Cleaved caspase-3 proteins in beta cells exposed to GLT for 12 h. (E) Proposed model of interactions between Mig6, NumbL & TRAF6 on EGFR and NFkB signaling in beta cells. *p< 0.05 vs. DMSO control under NG condition and #p<0.05 vs. DMSO control under GLT condition.

## 3. Discussion

In the present study, we discovered that Mig6 binds to NumbL under NG and this interaction is disrupted under GLT. Conversely, NumbL interacts more abundantly with TRAF6 under GLT than NG conditions. In addition, we established that NumbL down regulation prevents both the activation of NF-κB signaling and beta cell apoptosis under GLT. The combined interactions of Mig6, NumbL, and TRAF6, along with the pro-apoptotic nature of NumbL, may indicate that these proteins regulate beta cell apoptosis under GLT. Further, we propose a model that under NG conditions Mig6 is locked in an inactive complex with NumbL and thereby keeping pro-survival EGF signaling active. When beta cells are exposed to GLT conditions, this Mig6-NumbL complex is disrupted to release Mig6 and NumbL. The released Mig6 protein then binds to EGFR and inhibits EGF signaling, meanwhile NumbL binds to and activates TRAF6 protein to induce NF-κB signaling (**Figure 5E**). The results presented here are consistent with a model wherein GLT causes NumbL to switch from a Mig6-containing complex to a TRAF6-activating complex, thereby increasing pro-apoptotic signaling through NF-κB in beta cells (Figure 5E).

Given the known role for EGF signaling in pancreatic beta cell survival and proliferation [12, 17] and the ability of Mig6 to inhibit this pathway [40], we hypothesized that the Mig6 interacting partner NumbL would regulate EGF signaling. To our surprise, NumbL depletion significantly increased EGF signaling under NG and did not alter EGF signaling under GLT (**Supporting Information Figure S2B**). This increased EGF signaling under NG conditions can be partially be explained by the fact that NumbL, along with Numb protein, also participates in Erbb receptor (EGFR belongs to Erbb family of receptors) degradation as reported by others [34]. The observation that EGF signaling is increased under NG and impaired in GLT treated cells even after NumbL depletion indicates that the free Mig6 does not directly inhibit EGFR after release from Mig6-NumbL complex. We believe that the free Mig6 protein in NumbL depleted cells still needs to undergo posttranslational modifications before binding to EGFR and exerts its inhibitory function. Our group has observed that under GLT conditions, Mig6 is upregulated and is responsible for EGFR signal impairment (unpublished data). Consistent with this idea, other groups have reported that Mig6 undergoes extensive phosphorylation events before inhibiting EGF signaling [40]). Hence, further investigations are required to decipher the role of NumbL in EGF signaling in beta cells. It remains to be determined whether other factors, such as post-translational modification of Mig6, play a role in mediating the dynamic relationship between Mig6 and NumbL under GLT [40]. It also remains to be determined if the NumbL-Mig6 complex locks Mig6 in an inactive form and whether other pathways are involved in GLT-induced beta cell apoptosis. Bagnati *et al*. reported that under GLT, there is increased expression of CD40 receptors in beta cells [13] and CD40-TRAF6 interactions are known to activate the NF-κB pathway [13, 41]. Accumulation of NumbL-TRAF6 complex and initiation of NF-κB signaling under GLT suggests that NumbL plays a key role in activating NF-κB pathway. However, further investigations are warranted to understand how the TRAF6-NumbL complex activates the NF-κB pathway. We hypothesize that NumbL promotes polyubiquitination of TRAF6, which in turn activates the IKK complex. It also remains to be determined if CD40 receptor binding to this NumbL-TRAF6 complex is required for NF-κB activation.

TRAF6 activity depends on its polyubiquitination [16], and it is not clear if TRAF6, being an ubiquitin ligase, ubiquitinates itself or if other factors are involved [15, 16, 38]. Swarnker *et al*. determined that NumbL promotes polyubiquitination of TRAF6 and NEMO in osteoclasts [38]. As polyubiquitinated TRAF6 is required for activation of NF-kB signaling and NumbL promotes polyubiquitination of TRAF6, it is intuitive to predict that under GLT, NumbL activates TRAF6 to induce NF-kB and thereby causing beta cell apoptosis. However, Swarnker *et al*. reported NumbL negatively regulates NF-κB signaling in osteoclasts whereas we observe NumbL activating NF-κB signaling. The contradictions in these results can be explained partly by the difference in the context of the study, cell type and incubation time of the experiments. In our study, we treated beta cells with GLT for a short period of time (1 to 12 h) whereas Swarnker *et al*. used a longer duration cell model (4 d) to study the effect of NumbL on TRAF6. In beta cells, NF-κB activation starts within a couple of hours after exposure to GLT and apoptosis is induced at approximately 8 h exposure to GLT. Because polyubiquitinated TRAF6 first initiates the NF-κB pathway and then undergoes degradation, it is possible that we captured the initial phase of the events, such as TRAF6 ubiquitination, IkBa degradation and NF-κB activation, that might lead to apoptosis of beta cells. Further investigations are required to delineate these molecular events.

Overall, NumbL depletion seems to be beneficial for the survival of beta cells under diabetogenic conditions. To support this idea, we also observed that NumbL is significantly decreased in human Islets from prediabetic subjects, where there is expansion of functional beta cell mass. However, more investigations are warranted to validate NumbL as a drug target to treat diabetes.

### Conclusion

In summary, the results of the present study suggest that NumbL and Mig6 are in an inactive complex under NG conditions and under GLT; they separate and Mig6 decreases EGF signaling while NumbL increases NF-κB signaling. In addition, we uncovered a novel role of NumbL in beta cell apoptosis under GLT. Future investigations will evaluate NumbL as a potential therapeutic target to prevent beta cell death under diabetogenic conditions.

## 4. Methods

### 4.1 Materials

The following antibodies were purchased from Cell Signaling Technologies (Denver, CO, USA): phospho-EGFR, EGFR, p65, phosphor-p65, phosphor-eIF2a, eIF2a, Caspase-3 and Cleaved Caspase-3 (**Supporting Information Table S1**). Antibodies to NumbL and TRAF6 were obtained from SantaCruz Biotechnology (Dallas, TX, USA). The sequence of siRNAs (mission siRNAs purchased from Sigma-Aldrich, St. Louis, MO, USA) used against NumbL is: 5’- GAACUCACCUUUCAAACGU[dT][dT]-3’] & 5’- ACGUUUGAAAGGUGAGUUC[dT][dT]-3’ and Numb is: 5’- GAAGACUGAUUUCCCAAUA[dT][dT]-3’ & 5’- UAUUGGGAAAUCAGUCUUC[dT][dT]-3’; MISSION siRNA Universal Negative control was used as a control. Cell culture media, RPMI-1640 & CMRL-1066, were obtained from Thermo Fisher Scientific (Waltham, MA, USA). All the other reagents, if not indicated, were purchased from Sigma-Aldrich (St. Louis, MO, USA).

### 4.2 Human Islet cell culture

Cadaveric human islets were procured from the Southern California Islet Resource Center (City of Hope, Duarte, CA, USA) and cultured for 24 h in CMRL-1066 medium supplemented with 10% fetal bovine serum (FBS), 50 U/mL penicillin, and 50 ug/mL streptomycin before starting the GLT experiments. Then, the human islets were cultured in CMRL-1066 medium containing either 5 mM glucose (NG condition) or 25 mM glucose and 0.4 mM sodium palmitate (GLT condition) for various time points, as indicated in the figure legends. The islets were collected at the indicated time points and washed twice in PBS before cell lysis in RIPA buffer for immunoblot experiments. The protein lysates were loaded in a Wes instrument (ProteinSimple, CA, USA) according to the manufacturer’s protocol.

### 4.3 832/13 cell culture and transfection

Ins-1-derived 832/13 cells were cultured as described previously in Ins-1 medium, consisting of RPMI-1640 medium supplemented with 10% FBS, 50 U/mL penicillin, 50 *μ*g/mL streptomycin, 10 mM HEPES pH 8.0 buffer, 2 mM L-glutamine, 1 mM sodium pyruvate and 0.05 mM 2-mercaptoethanol [25]. The cells were seeded in 6-well tissue culture plates and, after overnight culture, were treated with 50 nM of either Mission control Negative siRNA (siControl) or siRNAs specific against Numb or NumbL (siNumbL) for 48 h. Then medium was replaced with either normal glucose (NG) or GLT media for the indicated time periods, as described in the figure legends.

### 4.4 Glucolipotoxicity experiments with 832/13 cells

832/13 cells were grown in Ins-1 medium until confluent and then cultured in Ins-1 medium containing 25 mM glucose & 0.4 mM palmitate for various time periods, as indicated in the figure legends. For NG conditions, the cells were incubated in Ins-1 medium containing 11 mM glucose.

### 4.5 Co-immunoprecipitation of Mig6 binding partners

832/13 cells were grown in 15 cm round tissue culture dishes to near confluence and then treated with adenoviruses expressing FLAG-tagged rat Mig6 (Ad-Flag-Mig6) or GFP (Ad-GFP) for 48 h. The cells were then treated either 5 mM glucose or 25 mM glucose and 0.4 mM palmitate for 12 h. After the incubation, the cells were collected and sonicated for 10 seconds with 60% amplitude in cell lysis buffer (20 mM Tris PH 7.2, 150 mM NaCl, 1% NP-40, 1 mM EDTA, 1 mM EGTA, 1 mM NaF, 1 mM sodium orthovanadate, 1 mM PMSF and Roche’s protease inhibitor tablet (Sigma-Aldrich, catalog # 4693159001). The cell lysate was centrifuged at 14,000 *xg* for 10 min to collect supernatant. Protein A/G sepharose beads were equilibrated with cell lysis buffer and incubated with the supernatant for 12 h at 4°C with rotation. After the incubation, the beads were collected by centrifugation at 1000 x*g* for 5 min and extensively washed in cell lysis buffer to remove non-specific binding proteins. The beads were then incubated with elution buffer (cell lysis buffer + 500 *μ*g/mL FLAG peptide) to specifically elute FLAG-tagged Mig6 and its binding partners. The eluate was then further processed and peptides were identified with mass spectrometric analysis by the Multi-Omics Mass Spectrometry Core at the City of Hope.

### 4.6 Apoptosis assay (cleaved caspase 3 activity assay)

832/13 cells were treated with either NG or GLT for the indicated time points. After the incubation, the cells were lysed in M-PER mammalian protein extraction reagent without protease inhibitors and the supernatant was collected after centrifugation at 14000 x*g* for 15 min. The protein concentration of the supernatant was determined by the BCA method. In a 96-well black microplate, 50 *μ*g of protein was incubated with 25 *μ*M of caspase 3 fluorometric substrate (Ac-Asp-Glu-Val-Asp-AMC) and the plate was read at 37°C for 1 h with excitation and emission wavelengths of 360 nm and 460 nm, respectively. In the same plate, a standard curve was prepared with AMC (7-Amino-4-methylcoumarin) to quantify the caspase 3 activity.

### 4.7 Proliferation assay (EdU incorporation)

Cell proliferation was quantified using EdU incorporation assay, as described before [26, 27]. Briefly, nearly 5,000 cells were seeded in 8-well chamber plates and treated with either siControl or siNumbL for 48 h and the incubated in GLT conditions for 12 h in the presence of 10 *μ*M EdU. The cells were then fixed with 4% formaldehyde for 30 min and permeabilized with 0.5% Triton X-100 for 15 min. The cells were then incubated for 30 min with detection reagent (100 mM Tris-Cl pH 8.5, 0.5 mM CuSO_4_, 25 *μ*M sulfo-Cyanine 3 azide and 50 mM ascorbic acid). After the incubation, the cells were washed in PBS and mounted using Vectashield mountant containing DAPI dye. Cell images were captured using a fluorescence microscope and the number of DAPI- and EdU-stained cells was counted using ImageJ software.

### 4.8 Immunoblotting

After treatment, cells were lysed in RIPA buffer and centrifuged at 14000 xg for 20 min to collect supernatant. Protein concentrations were determined by the BCA method and equal amounts of protein were run on SDS-PAGE gels. The proteins were then transferred to PVDF membranes and immunoblotted. The following primary antibodies with the dilution factor of 1: 1000 were used for immunoblotting: caspase-3, NumbL, Numb, p65, phospho-p65, IκBα, FLAG, TRAF6, γH2AX, phospho-eIF2a and total eIF2a. Tubulin and actin antibodies were used at 1:5000 dilutions. The following secondary antibodies were used: goat anti-mouse IgG-HRP conjugate (1:5000) & goat anti-rabbit IgG-HRP conjugate. The blots were developed using Amersham ECL detection reagent (GE Healthcare, Chicago, IL, USA). For samples derived from human islets, with a more limited amount of protein available, we performed the immune assay using the capillary Western-blot system (Wes, Protein Simple; San Jose, CA, USA), due to the sensitivity compared to the traditional western-blot method. All experimental steps were carried out according to the manufacturer’s instructions.

### 4.9 Glucose-stimulated insulin secretion (GSIS) assay

832/13 cells were treated with siControl or siNumbL as described above (section 2.3) and 48 h after treatment, a GSIS assay was performed as previously described [28, 29]. Briefly, cells were incubated in low glucose (2.5 mM) KRBB buffer for 1 h and then incubated in KRBB buffer containing low glucose (2.5 mM) or high glucose (15 mM) for 1 h. The cells and media were separated after the incubation and processed for quantification of insulin levels using the RI-13K Rat Insulin RIA Kit (Millipore) according to the manufacturer’s protocol.

### 4.10 Statistical analysis

All the quantified data are presented as means ± SEM. Student’s *t*-tests for analysis of two groups and ANOVA for analysis of more than two groups were employed to detect differences between groups. Tukey’s *post hoc* test was conducted after ANOVAs, and *p*-values of < 0.05 were considered statistically significant.

## Acknowledgements

This work was supported by a National Institutes of Health grant DK099311 to PTF and a Postdoctoral Fellowship Award from the American Heart Association to HDB. Human islets were generously provided by the Southern California Islet Resource Center. Nancy Linford, PhD, provided editing assistance. Research reported in this publication included work performed in the Comprehensive Metabolic Phenotyping and the Mass Spectrometry and Proteomics Cores at the City of Hope, with the latter supported by the National Cancer Institute of the National Institutes of Health under grant number P30CA033572. The content is solely the responsibility of the authors and does not necessarily represent the official views of the National Institutes of Health.

## Supporting Information

**Figure S1:**
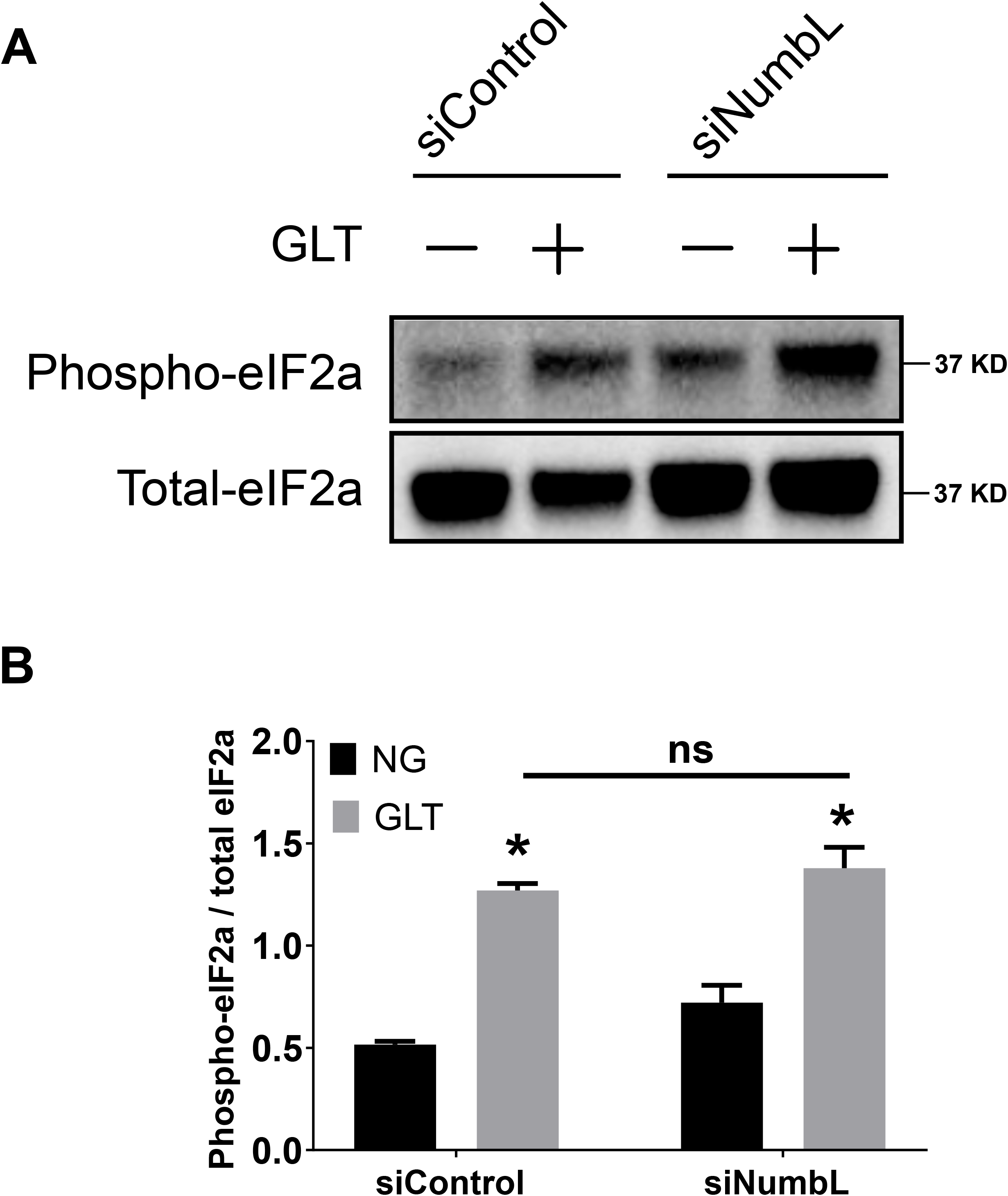
siRNA-mediated knock down of NumbL does not prevent GLT-induced ER-stress. (A) & (B) representative immunoblot and quantification of ER-stress marker proteins phospho-eIF2α and total eIF2α. 832/13 Ins1 cells were treated with NG or GLT conditions for 12 hours. *p<0.05 vs. siControl group under NG & ns; p>0.05.

**Figure S2:**
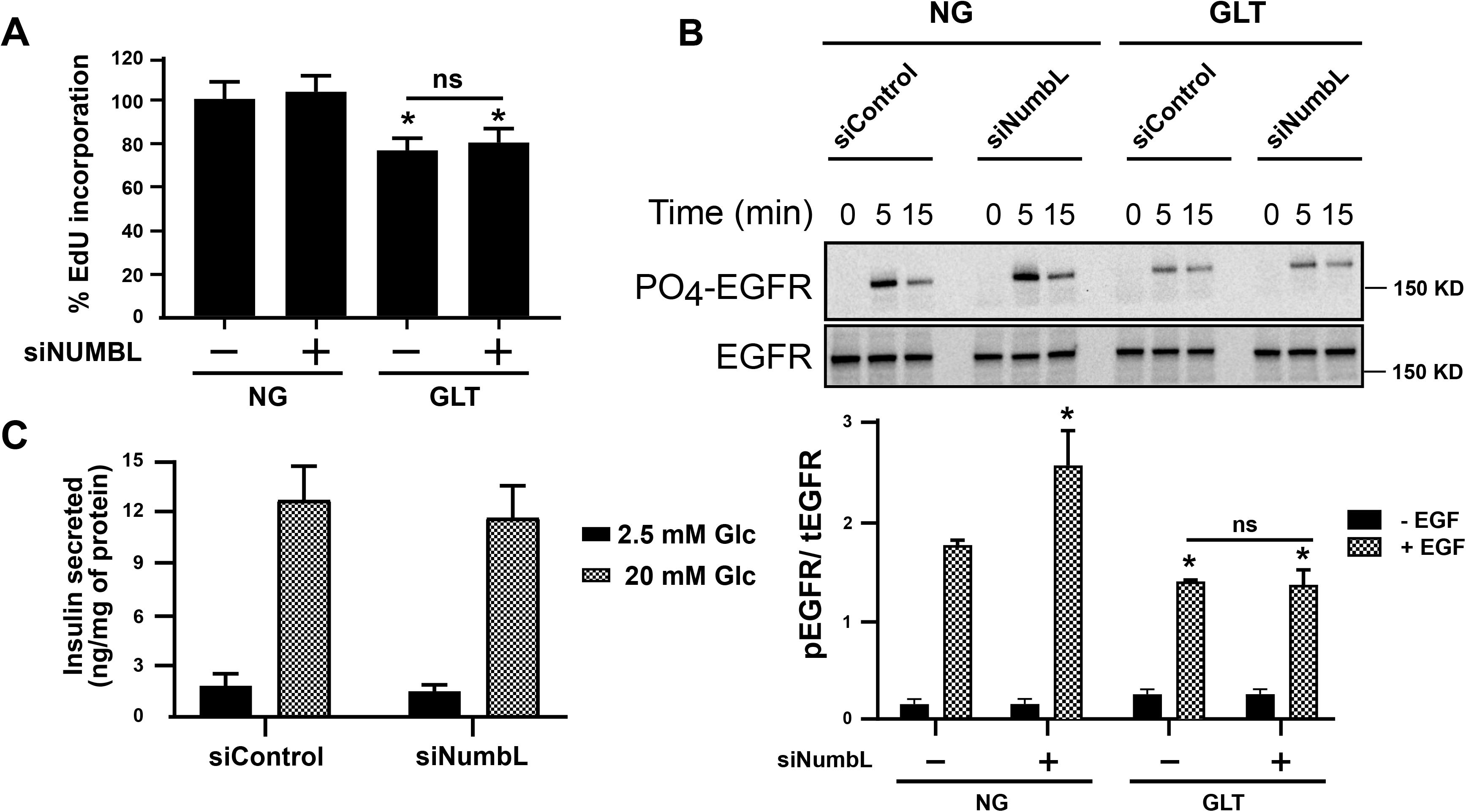
NumbL down regulation does not affect the proliferation and insulin secretion capacity of beta cells. (A) Quantification of beta cell proliferation using EdU incorporation assay in siNumbL- or siControl-treated cells under NG or GLT conditions for 12 h. (B) NumbL down regulation does not rescue impairment of EGF signaling under GLT. Representative immunoblot of phospho-EGFR and total EGFR proteins from siNumbL- or siControl-treated cells cultured under NG or GLT for 12 h. After incubation, the cells were exposed to either 0, 5 or 15 min of EGF ligand. Quantification of phospho-EGFR levels from EGF treated samples for 0 (-) and 5 min (+) are shown. *p<0.05 vs. siControl with 5 min EGF group under NG. (C) Quantification of glucose-stimulated Insulin secretion capacity of Ins1 832/13 cells in siNumbL- or siControl-treated cells. *p<0.05 vs. siControl group under NG & ns; p>0.05.

**Figure S3:**
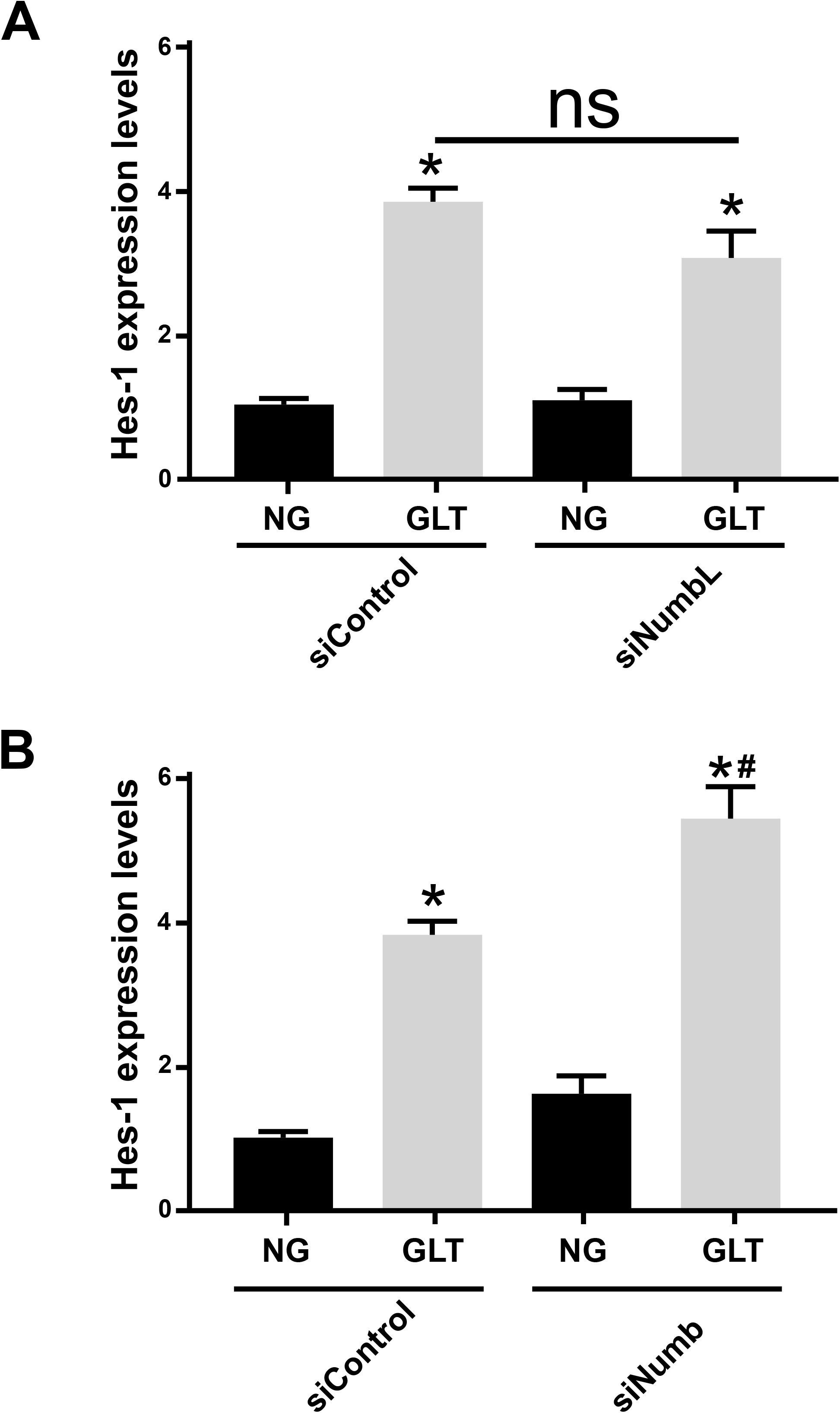
NumbL down regulation does not affect Notch signaling in beta cells. (A) Quantification of Hes-1 mRNA levels in siNumbL- or siControl-treated cells. (B) Quantification of Hes-1 mRNA levels in siNumb or siControl treated cells. Cells were treated with NG or GLT conditions for 12 h. *p<0.05 vs. siControl group under NG & ns; p>0.05.

**Supporting Information - Table 1:** The details of the antibodies used in the manuscript:

| **Antibody** | **Company** | **Catalog #** | **Dilution** |
| --- | --- | --- | --- |
| Phospho-EGFR (Y1173) | Invitrogen | 44794G | 1:1000 |
| EGFR | Cell Signaling Technology | 2232 | 1:1000 |
| NUMB | Cell Signaling Technology | 2756 | 1:1000 |
| NUMBL | Santa Cruz Biotechnology | SC-390590 | 1:1000 |
| Phospho-p65 | Cell Signaling Technology | 3031 | 1:1000 |
| p65 | Cell Signaling Technology | 8242 | 1:1000 |
| Phospho-eIF2a (S51) | Cell Signaling Technology | 9721 | 1:1000 |
| eIF2a | Cell Signaling Technology | 5324 | 1:1000 |
| Caspase-3 | Cell Signaling Technology | 9662 | 1:1000 |
| Cleaved Caspase-3 | Cell Signaling Technology | 9664 | 1:1000 |
| TRAF6 | Santa Cruz Biotechnology | SC-8409 | 1:1000 |
| Tubulin | Cell Signaling Technology | 2144 | 1:1000 |
| Actin | Millipore Sigma | MAB1501 | 1:1000 |

